# Alpaca: a kmer-based approach for investigating mosaic structures in microbial genomes

**DOI:** 10.1101/551234

**Authors:** Alex N. Salazar, Thomas Abeel

## Abstract

**Summary:** Microbial genomes are often mosaic: different regions can possess different evolutionary origins due to genetic recombination. The recent feasibility to assemble microbial genomes completely and the availability of sequencing data for complete microbial populations, means that researchers can now investigate the potentially rich evolutionary history of a microbe at a much higher resolution. Here, we present Alpaca: a method to investigate mosaicism in microbial genomes based on kmer similarity of large sequencing datasets. Alpaca partitions a given assembly into various sub-regions and compares their similarity across a population of genomes. The result is a high-resolution map of an entire genome and the most similar scoring clades across the given population.

**Availability:** https://github.com/AbeelLab/Alpaca

## 1 Introduction

The ever-increasing availability of genomic data is enabling researchers to obtain novel insights about the genetic diversity and evolutionary history of various organisms. This is particularly important for microbes as they can originate from widely-diverse populations due lateral exchange of genetic information and hybridization of multiple genomes (Ochman *et al.*, 2000; Rainieri *et al.*, 2006). For example, horizontal gene transfer can lead to the introgression of novel sequences in nuclear chromosomes, aiding the adaptation for certain environments (Ochman *et al.*, 2000). The resulting genome may thus be mosaic, meaning that different genomic segments may possess different evolutionary origins (Martin, 1999). Genome hybridization—the joining of two or more genomes from different strains/species—can also lead to mosaic structures due to recombination of homologous segments followed by selection of those segments with fitness advantages (Dujon and Louis, 2017). As such, the genome of an individual microbe may be the result of a rich history of genetic adaptations after coming in contact with different populations (Martin, 1999; Ochman *et al.*, 2000; Ochman *et al.*, 2000; Dujon and Louis, 2017). In other words, there can be multiple origins for the genetic content in a single genome.

We are currently at a crossroads of microbial genome sequencing data-types: single-molecule sequencing technologies (such as PacBio and Oxford Nanopore) are enabling complete characterization of (individual) microbial genomes (Loman *et al.*, 2015; Rodríguez *et al.*, 2015; Salazar *et al.*, 2017; Yue *et al.*, 2017), while past and current short-read sequencing projects of hundreds to thousands of microbial strains contain vast information about their population diversity (Manson *et al.*, 2017; Peter *et al.*, 2018). Combining both genomic data-types can thus aid in our understanding of the potential mosaic structures in microbial populations. Here, we present Alpaca: a stand-alone method for investigating mosaic structures in microbial genomes.

## 2 Alpaca description

Given a reference genome assembly and a set of read-alignments, Alpaca will identify the most similar sample(s) in the population for various subregions across the genome based on kmer similarity. This is done in three major steps: (i) partitioning a genome into subregions, (ii) scoring the similarity of each subregion for every sample in the population, and (iii) a summary of all subregions with their top-scoring sample(s); ultimately providing insights about the genetic origins and potential mosaic structures of a genome.

We provide a user-friendly, command-line implementation of these steps along with high-resolution and informative visualizations. Alpaca is written in the Scala programming language (https://www.scala-lang.org/) and packaged with a stand-alone binary distribution. Alpaca greatly benefits from parallelization (i.e. availability of multiple CPUs and multiple chromosome sequences) as it is implemented under a functional paradigm. We describe the major features of Alpaca below.

### 2.1 Alpaca database: genome partitioning of subregions

The first step is to create an *Alpaca database* for a given reference genome. The input is a FASTA-formatted assembly, a sorted BAM file of the native read alignments to the assembly (i.e. the same reads used to create the assembly), and the estimated genome size. Alpaca will then iterate through each FASTA-entry and create non-overlapping subregions. Unique kmers are extracted for each subregion sampling from both the assembly and read-alignments independently. To minimize erroneous kmers, users can specify a minimum kmer count or allow Alpaca to automatically detect the threshold based on alignment coverage. The output is a sub-directory describing all subregions and their corresponding kmer-sets.

### 2.2 Target comparison: computing subregion similarity

The next step is to compute the similarity of all subregions in the reference genome against a target genome. The input is the path to the Alpaca database of the reference genome (see above), read alignments to the reference-genome’s assembly as a sorted BAM file, and the estimated genome size of the target genome. Alpaca will then iterate through all subregions in the database and construct kmer-sets for the target genome using the same procedure in the sub-section before. A similarity score for a given subregion can thus be computed as the Jaccard Index of the two corresponding kmer-sets:

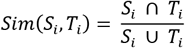

Where *S*_*i*_ corresponds to the kmer-set from the reference genome and *T*_*i*_ the kmer-set from the target genome. The output is a tab-delimited file containing the coordinate of every sub-region in the database and the similarity score.

### 2.3 Population summary: ranking top-scoring samples

Comparing the subregion similarities against a set of target genomes can provide insights for potential mosaic structures in the reference genome. The final step is thus to summarize every similarity score by ranking and retaining the top scoring target sample(s) for each subregion. The input is the path to the Alpaca database of the subject genome and a tab-delimited file listing the path of the target comparison outputs from the previous step (see above). Alpaca will then iterate through each subregion and retain the top-scoring targets. Note that there may be multiple top-scoring samples since different samples may possess the same similarity score. Alpaca will only retain the scores of subregions possessing a (configurable) number of top-scoring samples. The remaining samples have their similarity value set to zero, i.e. no similarity, to mitigate ambiguous subregions.

The output is a tab-delimited file of every subregion along with its top-scoring sample(s).

#### 2.3.1 Alpaca-layout: visualizing mosaic genomes

If the used target population possesses labels (i.e. lineages, clades or species), then a high-resolution visualization for interpreting mosaic structures can be created (see Figure 1A). Using the population summary file, Alpaca will iterate through each chromosome and draw it as a sequence of rectangles representing individual subregions across the chromosome (see Figure 1A). The color of each subregion represents the corresponding label of the top-scoring sample(s). Note that there can be multiple colors for each subregion, whose proportions are based on the number of labels in the top-scoring samples. The corresponding proportion of a color for each subregion is computed as the similarity score, *sim*(*S*_*i*_, *T*_*i*_), multiplied by the proportion of samples belonging to each label. To display *unexplained* similarity (i.e. sub-regions that are ambiguous or have low-scoring similarities), 1 − *sim*(*S*_*i*_, *T*_*i*_) is drawn as white.

**Figure 1.**
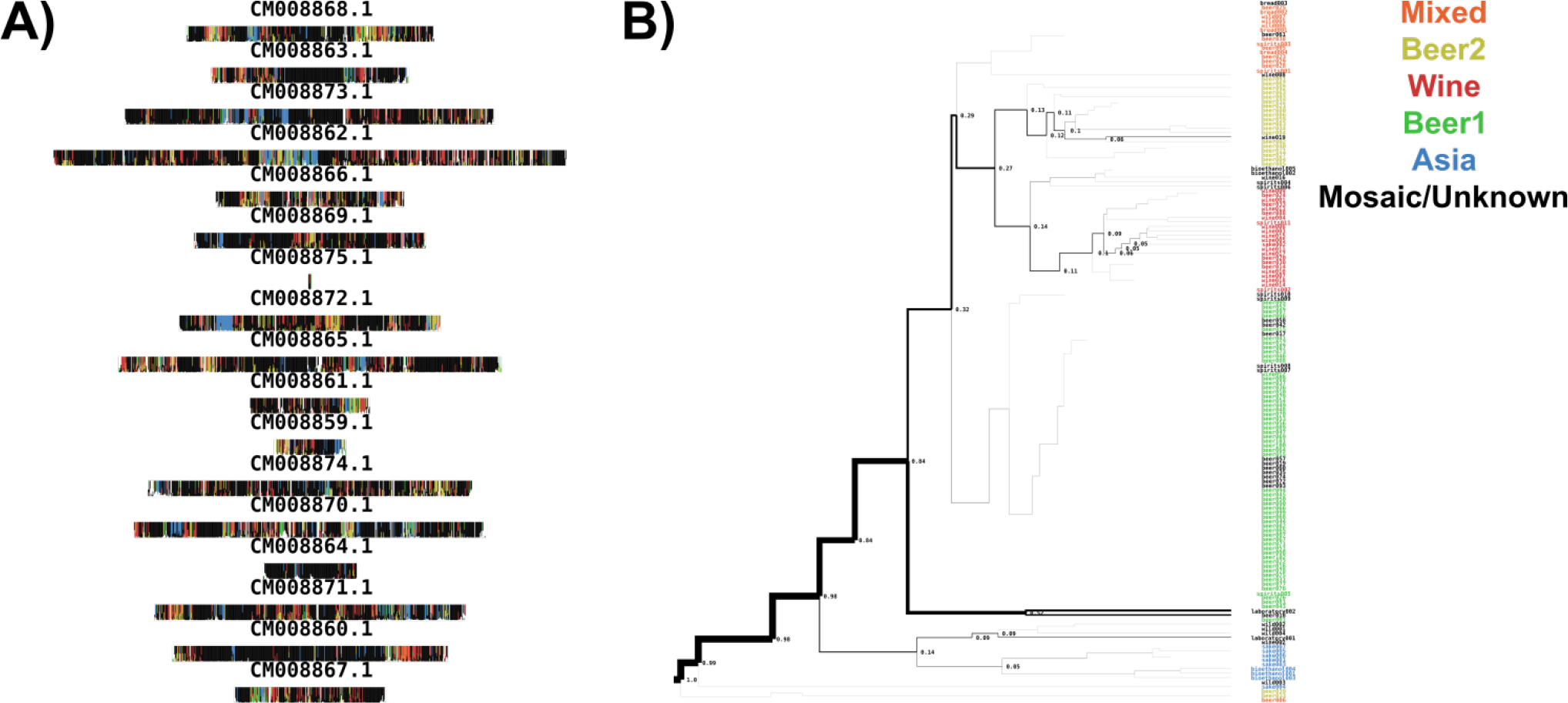
Summary visualizations of the mosaic profiles of the *S. cerevisiae* strain, CEN.PK113-7D, using 155 *S. cerevisiae* strains from Gallone *et al*. (A) Alpaca-layout figure: each rectangle is a chromosome from the assembly which is composed of a sequence of smaller rectangles (subregions) whose colors are based on the lineages (see colored legend in Fig 1B) from the top-scoring samples within that subregion. (B) Tree-tracing figure: the tree is a hierarchical clustering of the 155 strains and the width of the branches correspond to the overall frequency for which a given strain was a top scoring sample for any subregion. The color of the strain corresponds to the evolutionary lineage (Mixed, Beer2, Wine, Beer1, Asia, Mosaic/Unknown) as defined by Gallone *et al.*

The resulting visualization (SVG-formatted) displays every subregion and the corresponding similarity to assigned lineages or species, enabling users to identify instances of mosaic structures.

#### 2.3.2 Tree-tracing: similarity across population-structures

The Alpaca-layout visualization is in context of lineages or species and therefore does not provide information about subregion similarity to individual samples or sub-populations. If a (phylogenetic) tree in Newick-format (describing the population structure of different lineages or species) is provided, Alpaca will traverse the tree and hierarchically display the frequency of top-scoring samples and their corresponding (sub-)clades (see Figure 1B). Starting at the root-node, Alpaca will traverse through the tree and draw each branch with a thickness corresponding to the proportion of a current node’s children as top-scoring samples over the total sum.

## 3 Runtime and conclusion

We applied Alpaca to investigate potential mosaic profiles of the industrial *S. cerevisiae* strain, CEN.PK113-7D. In general, the database creation took ~1.5 min with less than 1 gb of ram requiring ~57 mb of space with a single CPU using default parameters. We aligned short-read Illumina data from 155 *S. cerevisiae* strains from Gallone *et al.* to the CEN.PK113-7D long-read assembly using BWA and computed genome similarities with Alpaca using default parameters. On average, target comparisons took ~2.5 hrs with less than 2 gb of ram on two CPUs. The results are summarized in Figure 1.

To conclude, Alpaca can provide further insights of microbial assemblies by characterizing potential mosaic structures across sequencing datasets of microbial populations.

## Acknowledgements

We would like to thank Arthur R. Gorter de Vries, Jean-Marc Daran, and Niels Kuipers for critical discussion.

## Funding

This research was funded by a grant from BE Basic Foundation related to FES funds from the Dutch Ministry of Economic Affairs.

## Conflict of Interest

none declared.

